# Canonical Wnt signalling is activated during BEC-to-hepatocyte conversion *in vivo* and modulates liver epithelial cell plasticity in hepatic organoids

**DOI:** 10.1101/2020.11.09.374462

**Authors:** Eider Valle-Encinas, Niya Aleksieva, Carmen Velasco Martinez, Michael Dawes, Matthew Zverev, Miryam Müller, Anika Offergeld, Matthew J Smalley, Catherine Hogan, Stuart J Forbes, Tom G Bird, Trevor C Dale

## Abstract

While it is recognized that the Wnt/β-catenin pathway orchestrates hepatocyte proliferation in both homeostasis and injury, little is known about the importance of β-catenin in biliary epithelial cell (BEC) plasticity. In this study, the dynamics of activation of the canonical Wnt pathway were investigated during BEC-to-hepatocyte conversion using as a model methionine/choline deficient (MCD)-injured livers where hepatocyte proliferation was compromised by the overexpression of p21. In this model, activation of β-catenin was found an event associated with BEC reprogramming. Using ductal organoids to model BECs transitioning into hepatocytes, we found that activation of the Wnt/β-catenin pathway in these cells promoted partial escape from a biliary fate and triggered the acquisition of progenitor cell features. Our data furthermore support that BECs are source of Wnt ligands and that Rspo proteins potentially act as the limiting factor controlling the activation of β-catenin activation and BEC reprogramming during severe liver damage.

## Introduction

The Wnt pathway plays a central role in the development, homeostasis and regeneration of the liver (Perugorria et al., 2018; Si-Tayeb et al., 2010). Activation of the canonical branch in the pathway is initiated by the binding of Wnt ligands to two receptors, a member of the Frizzled family (Fzd1-10) and a receptor from the Lrp family (Lrp5-6), culminating in the stabilization of the transcriptional co-activator β-catenin. In the unperturbed adult liver, transcriptional activation by β-catenin is restricted to pericentral hepatocytes, where it regulates the expression of metabolic genes ((Ben-Moshe and Itzkovitz, 2019; Burke and Tosh, 2006; Cadoret et al., 2002; Gougelet et al., 2014). The Wnt pathway has additionally been associated with the regenerative responses of hepatic progenitor cells (HPCs) that arise upon chronic liver injury (Apte et al., 2007; M. Hu et al., 2007; Huch et al., 2013; Itoh et al., 2009; Lin et al., 2017; Zhang et al., 2008). The identity and origin of HPCs has been a topic of interest for many years and has recently been studied in mice using a range of different model systems that differ in injury mechanism and the degree to which rescue from the hepatocyte compartment is inhibited (Deng et al., 2018; Perugorria et al., 2018; Raven et al., 2017; Russell et al., 2019). As our knowledge in the different hepatobiliary regenerative strategies of the liver expands, the exact role of Wnt pathway in diverse pathophysiological contexts needs to be clarified.

The liver harbours two epithelial cell populations: hepatocytes and biliary epithelial cells (BECs) with a common epithelial progenitor; the hepatoblast. Upon injury, the repair strategy of both epithelial compartments greatly depends on the type of injury and the extent of the damage (Ng and Ananthakrishnan, 2018; Monga, 2020). In mild/moderate liver injury, hepatocytes (Hnf4a+) are ‘democratic’ and surviving healthy cells proliferate and regenerate the damaged parenchyma (Chen, 2020; Matsumoto et al., 2020; Monga, 2020; Raven et al., 2017; Deng et al., 2018; Schaub et al., 2018; Tarlow et al., 2014; Yanger et al., 2013). This process is orchestrated by canonical Wnt ligands and Wnt agonists such are members of the Rspo family that are upregulated in response to liver injury (Planas-Paz et al., 2019; Rocha et al., 2015; Sun et al., 2020). Similarly, the regeneration of the biliary epithelium is fuelled by cells committed to the BEC lineage (Krt19+, Hnf1b+ cells) (Jörs et al., 2015; Raven et al., 2017). While initial studies suggested that BEC regeneration was dependent on canonical Wnt signals, recent studies failed to identify activation of the Wnt/β-catenin pathway in the biliary epithelium of DDC injured animals (Apte et al., 2007; M. Hu et al., 2007; Huch et al., 2013; Lin et al., 2017; Pepe-Mooney et al., 2019; Planas-Paz et al., 2019; Wilson et al., 2020). Furthermore, concomitant loss of the canonical Lrp5/6 receptors (Lrp5/6^fl/fl^; Alb-Cre) or the Rspo receptors Lgr4 and Lgr5 (using Lgr4/5^fl/fl^; Alb-Cre) in the biliary epithelium did not cause a decline in the number of proliferative BECs nor the formation of A6-positive ductular structures in DDC-injured livers, suggesting that proliferation of BECs following injury occurs in a β-catenin - independent manner (Okabe et al., 2016; Planas-Paz et al., 2019).

Hepatocytes and BECs demonstrate a remarkable capacity for lineage plasticity when damage is prolonged or when the proliferation of the initial cell type is compromised. BEC-to-hepatocyte conversion has been exclusively reported when the proliferation of the pre-existing hepatocytes is impaired, which in murine models may be achieved by the overexpression of p21, genetic ablation of β1-integrin or β-catenin, or prolonged exposure to hepatotoxic insults (Deng et al., 2018; Manco et al., 2019; Raven et al., 2017; Russell et al., 2019). HPCs that were first described as cells that arise in response to injury with hepatocytes/BEC intermediate features have recently been suggested to be derived from ‘activated’ BECs and hepatocytes that partially escape from their committed state. The appearance of HPCs is thought to be associated with the creation of cellular niches in response to chronic damage and regeneration such as an unbalanced diet or viral infection. BEC-to-hepatocyte conversion in particular may be of special clinical relevance in liver pathological settings resulting from chronic and repeated injury such is long-term alcohol abuse (Aguilar Bravo et al., 2019).

While BEC proliferation during biliary repair has been shown to be independent of canonical Wnt signalling, the relative contribution of the Wnt/β-catenin pathway to BEC-to-hepatocyte conversion remains largely unexplored (Pepe-Mooney et al., 2019; Planas-Paz et al., 2019). In this study, we explored the role of canonical Wnt signalling in BEC cellular plasticity. Using a methionine/choline deficient (MCD) diet liver injury model in combination with p21 overexpression, we show that BEC-derived cells respond to canonical Wnt signals when hepatocellular regeneration is compromised. Using differentiated bile duct-derived organoids (BD organoids), we provide evidence that activation of the Wnt/β-catenin pathway in BECs enables a partial escape from their biliary-committed state and promotes the acquisition of a progenitor cell status. Altogether, our data supports a role for the canonical Wnt pathway in BEC plasticity.

## Results

### Axin2, Lrp6 and Lgr5 are dynamically expressed during BEC-to-hepatocyte conversion in the MCD p21 mouse model

To examine the dynamics of canonical Wnt activation during BEC reprogramming, we utilized a model of published by Raven et al. (2017) in which hepatocyte proliferation is prevented by the expression of p21 and BECs repopulate the hepatocellular parenchyma following MCD/recovery injury diet regime (Raven et al., 2017). In this model, BEC cells were lineage-traced by tamoxifen-induced labelling of Krt19 expression cells with tdTom (Krt19-CreERT2+/-; R26-LSL-tdTom+/+) and the biliary epithelium was labelled with tamoxifen with a ~40% recombination efficiency (Figure1, A) (Raven et al., 2017). As expected, BEC-derived hepatocytes (HNF4a+ tdTom+ cells) were only detected in MCD-injured animals overexpressing p21 (AAV8.p21) but not in the control group that received the AAV8 empty backbone control vector (AAV8.null) (Figure1, B).

**Figure 1.**
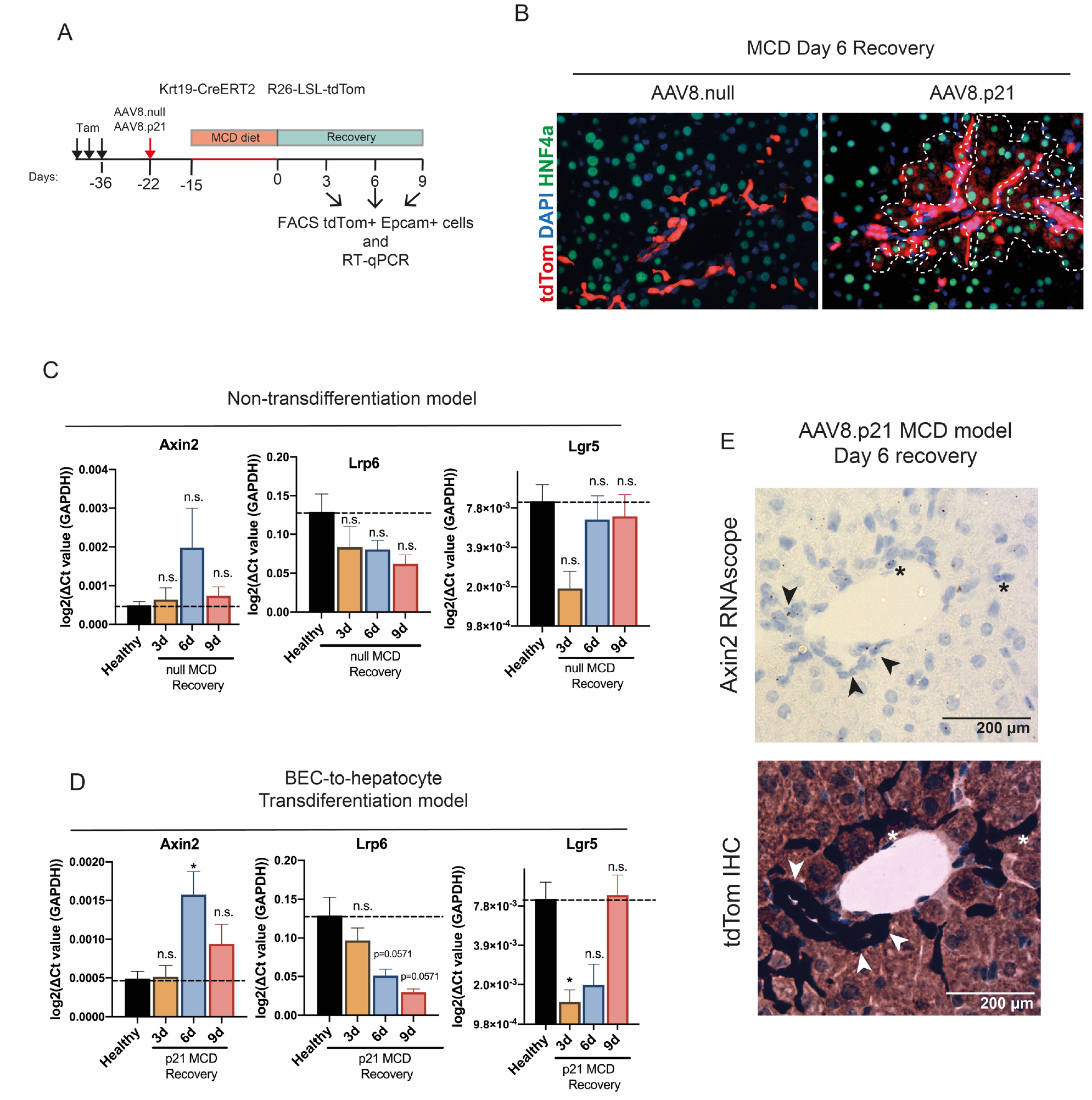
(A) Treatment time for the flow cytometry isolation of BECs (tdTom+ Epcam+ cells) from MCD-injured livers following administration of either AAV8.p21 or AAV8.null viral particles via tail injection. (B) Representative images of MCD-injured livers following 6-day recovery. Dash line highlights the presence of HNF4a tdTomBEC-derived hepatocytes. (n=4 animals) (C and D) RT-qPCR gene expression analysis of tdTom+ Epcam+ cells isolated at various time points from the AAV8.null (non-transdifferentiation, panel C) and the AAV8.p21 (BEC-to-hepatocyte transdifferentiation, panel D) MCD injury models. (n=4 animals per condition). (E) Axin2 RNAscope analysis of serial sections of AAV8.p21 MCD-injured livers at day recovery 6 (top). Immunohistochemistry for tdTom labels cells of ductal origin (bottom). Arrowheads point to BECs in ducts. Asterisks point to BECs Axin2+ cells showing an invasive phenotype. (n=3 animals).

To focus on BEC-derived cells prior to their adoption of an overt hepatocyte morphology and gene expression program, recombined EpCAM+ BECs positive for tdTom expression were isolated by FACS from the MCD AAV8.p21 and AAV8.null injury models for RT-qPCR analysis (Figure S2). Expression of the Wnt targets Axin2, and the Wnt pathway receptors / targets Lrp6 and Lgr5 was examined at recovery days 3, 6 and 9 (Figure1, A). Similar patterns were observed in tdTom+ EpCAM+ cells from both control and AAV8.p21-injected animals, although the effect size was greater in the later (Figure1, C and D). The expression of Lrp6 gradually decreased during recovery (Figure1, D), while the expression of the Rspo receptor Lgr5 recovered to levels present in uninjured animals within 9 days of recovery. Axin2 was maximally expressed on day 6 of recovery in the biliary epithelium of AAV8.p21 MCD-injured livers. Axin2 transcripts were also detectable in tdTom+ bile ducts and in BECs with an invasive phenotype by RNAscope in AAV8.p21 MCD-injured livers at day 6 of recovery (Figure1, E). Taken together, these data suggest that the canonical Wnt pathway is activated in the biliary epithelium of MCD-injured livers when hepatocellular regeneration is impaired.

### Differentiated BD organoids mirror aspects of BEC-to-hepatocyte conversion *in vitro*

To interrogate the role of the Wnt/β-catenin pathway in BEC-to-hepatocyte conversion, we moved to an *in vitro* system described by Huch et al. (2013) in which activated BECs grown as organoids transition into hepatocyte-like cells in culture (Huch et al., 2013). As expected, the expression of HPC-related markers (Tert, Cd44, Trop-2 and Lgr5) in BD organoids was decreased following 13-days of differentiation while the expression of hepatocyte lineage (Cebpa, Hnf4a and Prox1) was increased (Figure 2, A-C). In line with previous data, BD organoids increased the expression of hepatocyte metabolic markers (Alb, Fah, Ass1, Cyp2f2, Cyp2e1 and Cyp1a2) following differentiation although the transcript levels of these markers remained ~13-1200 fold lower than that observed in pooled primary hepatocytes (Figure2, D) (H. Hu et al., 2018; Huch et al., 2013). These results indicate that that while the differentiation conditions of Huch et al. (2013) were permissive for the commitment of cells to the hepatocyte lineage, organoid cells did not fully acquire a mature hepatocyte state in culture. Deng et al. (2018) described BECs, in their conversion to hepatocyte fate, acquire a bi-phenotypic state in which they co-express hepatocyte (HNF4a) and BEC (CK19) lineage markers *in vivo* (Deng et al., 2018). IF analysis of the differentiated organoids revealed co-expression of both HNF4a and CK19 markers (Figure 2, E), further supporting that BD organoids do not accurately reflect a fully matured hepatocyte phenotype and that these structures capture aspects of the biology of activated BECs transitioning into hepatocyte fate.

**Figure 2.**
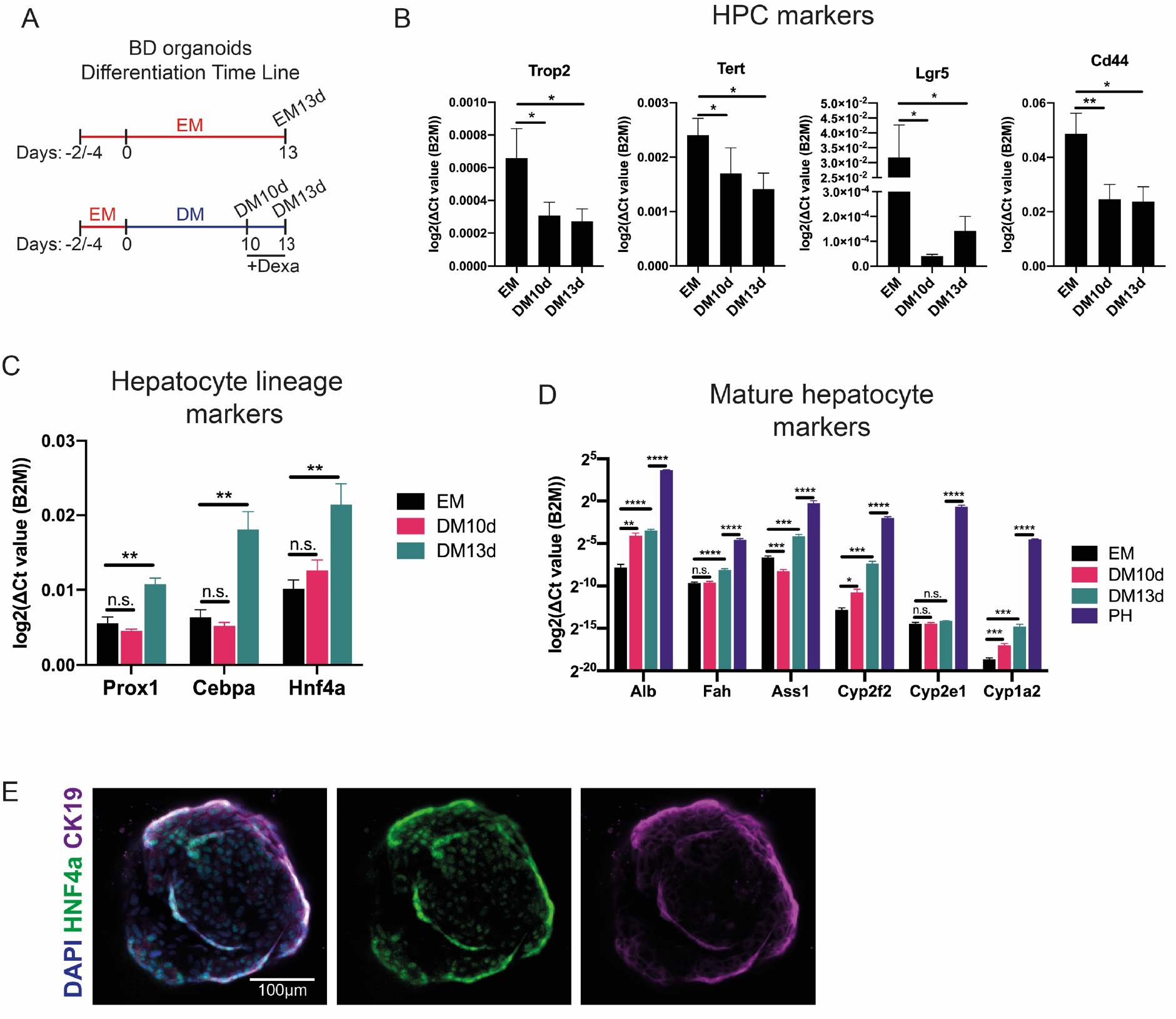
(A) BD organoids differentiation time line. Expansion medium (EM); differentiation medium day 10 (DM 10d); differentiation medium day 13 (DM 13d). (B-D) RT-qPCR gene expression analysis of HPCs (panel B), hepatocyte lineage (panel C) and hepatocyte metabolic markers (panel D) in BD bile duct organoids upon differentiation. For organoid samples, n=9 independent replicates (3 organoid lines were derived from 3 different wild-type animals. Each organoid line was differentiated using a different differentiation batch. 3 organoid wells were cultured per organoid line and processed independently (Table S2)). For primary hepatocyte samples, n=3 animals. SEM error bars. Statistics were determined using paired t-test (panels B and C) and multiple t-test (panel D). (E) Representative IF images show that BD organoids co-express the bile duct marker CK19 (magenta) and the nuclear hepatocyte lineage marker HNF4a (green) at day 11 of differentiation. Cell nuclei were counterstained with DAPI (blue).

### Activation of the Wnt signalling pathway in differentiated BD organoids drives the expression of HPC/proliferation genes

Having characterised the BD organoid system, we derived an organoid line from a Tet-O-ΔN89β-catenin mouse model (Jardé et al., 2012). In this organoid line, exposure to 0.1 mg/ml Dox during the last 3 days of differentiation induced expression of a non-degradable active mutant form of β-catenin with no evidence of toxicity (Figure S3). Stabilization of β-catenin (0.1 mg/ml Dox) in BD organoids resulted in the induction of canonical Wnt targets (Axin2 and Lgr5) and the HPC markers (Sox9, Spp1 and Cd44) (Figure 3, A and B). Activation of the pathway at the receptor level with Rspo1 (100 ng/ml) resulted into a more pronounced induction of these genes (Figure3, B). The Rspo1-HPC driven signature was lost in the presence of LGK974 (500 nM), a porcupine inhibitor that blocks secretion of Wnt ligands. This effect that was attenuated when exogenous Wnt3a was added to the media, suggesting that the induction of HPC genes by Rspo1-required the presence of endogenous Wnt ligands (Figure3, C). Rspo1 treatment also caused a significant increase in the expression of the proliferation markers Ki67 and CyclinD1 (Figure3, B). Exposure to either Dox or Rspo1 reduced the expression of both hepatocyte (Hnf4a) and BEC (Krt19) lineage markers, suggesting that activation of the canonical Wnt pathway induces escape from biliary fate and prevents the progression towards hepatocyte fate (Figure3, D). Of particular note, neither of the Wnt activation treatments induced the expression pericentral metabolic genes such as GS (Glul), Cyp1a2 and Cyp2e1, a response previously reported in hepatocytes both *in vivo* and *in vitro* (Figure3, E) (Gougelet et al., 2014; Wahlicht et al., 2020). These results further indicate that BD organoids are structures unable to recapitulate mature hepatocyte biology.

**Figure 3.**
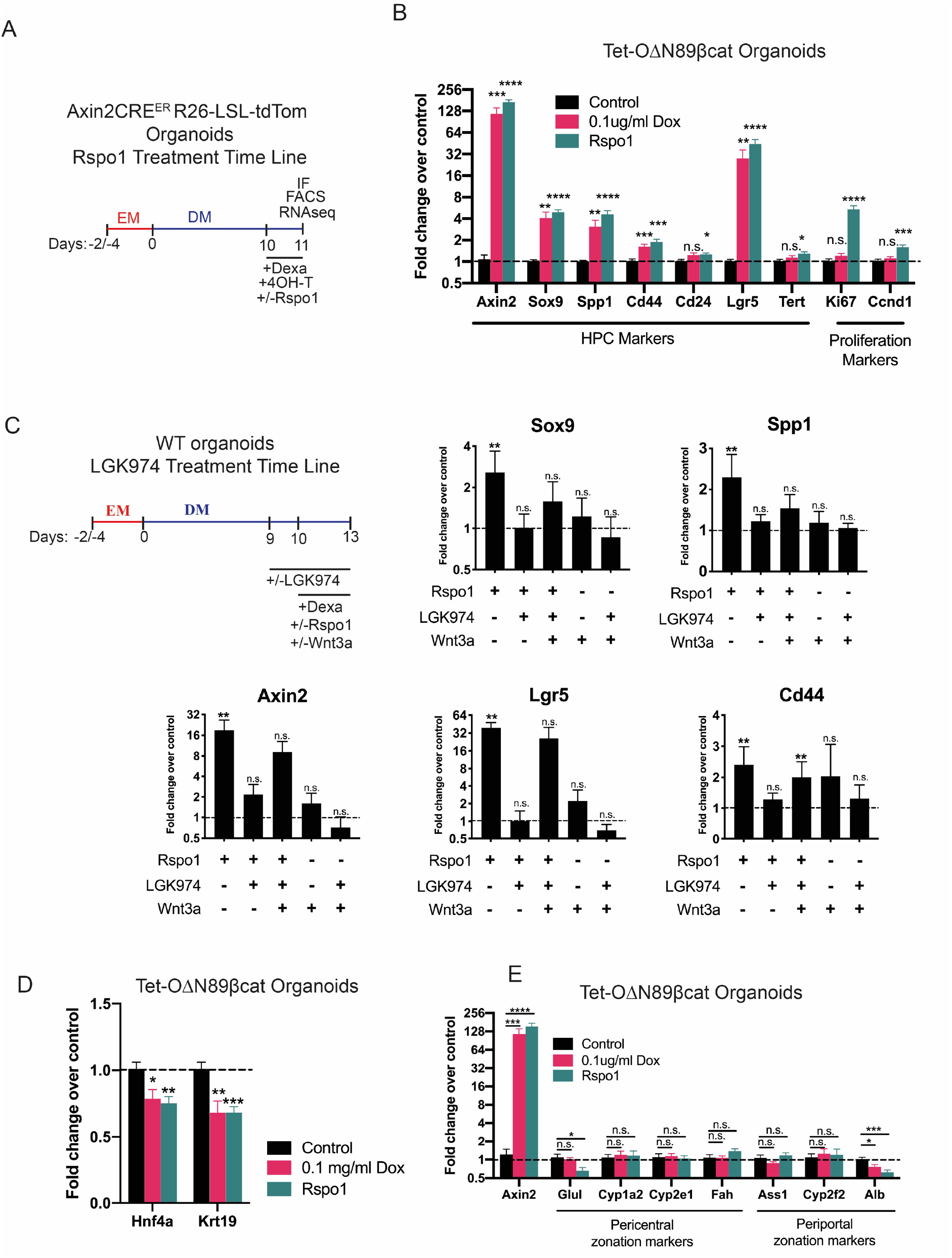
(A) Treatment timeline for Tet-OΔN89βcat BD organoids (panels B, D and E). (B) RT-qPCR gene expression analysis in Tet-OΔN89βcat BD organoids upon activation of mutant β-catenin (Dox) or exposure to Rspo1. n=9 (organoids derived from a single animal were analysed at 3 different passages using 3 different differentiation batches. In each passage, organoids were cultured in 3 wells that were processed independently (Table S2)). SEM error bars. Statistics were determined using multiple t-test. (C) RT-qPCR gene expression analysis of wild type organoids shows that exposure to LGK974 blocks Rspo1 induction of stem cell genes. LGK974 effects were partially rescued by the addition of exogenous Wnt3a. As LGK974 vehicle conditions, organoids were exposed to the equivalent DMSO concentration. n=5 (organoids derived from 4 wild type animals were analysed at 5 different passages using 4 different differentiation batches (Table S2)). SEM error bars. Statistics were determined using Mann-Whitney test. Dash line indicates gene expression levels in control organoids differentiated in the absence of LGK974, Rspo1 and Wnt3a. (D-E) RT-qPCR gene expression analysis in Tet-OΔN89βcat organoids upon activation of mutant β-catenin (Dox) or exposure to Rspo1 (see panel A). n=9 (organoids derived from a single animal were analysed at 3 different passages using 3 different differentiation batches. In each passage, organoids were cultured in 3 wells that were processed independently (Table S2)). SEM error bars. Statistics were determined using multiple t-test.

### Exposure of differentiated BD organoids to Rspo1 induces the appearance of an Axin2 expressing population with progenitor features

Axin2 expression was increased in the biliary epithelium during BEC-to-hepatocyte conversion *in vivo* (AAV8.p21 MCD injury model). To explore whether cells could be distinguished based on Axin2 expression in the organoid system, a line of BD organoids was established from an Axin2CreERT2+/-;R26-LSL-tdTom+/- transgenic mouse model in which Axin2 expressing cells are lineage traced as tdTom+ cells following exposure to 4-hydroxy tamoxifen (4-OHT). Cells expressing tdTom+ were rarely found (<0.4%) in differentiated Axin2CreERT2+/-;R26-LSL-tdTom+/- organoids exposed to 4-OHT for 24h both by flow cytometry and IF (Figure 4, A-C). By contrast, 4-10% of the total cells were labelled with tdTom+ when Rspo1 was concurrently added with 4-OHT for the last 24h of culture, thus confirming our previous observations of Axin2 upregulation following Rspo1 treatment (Figure 4, A-C).

**Figure 4.**
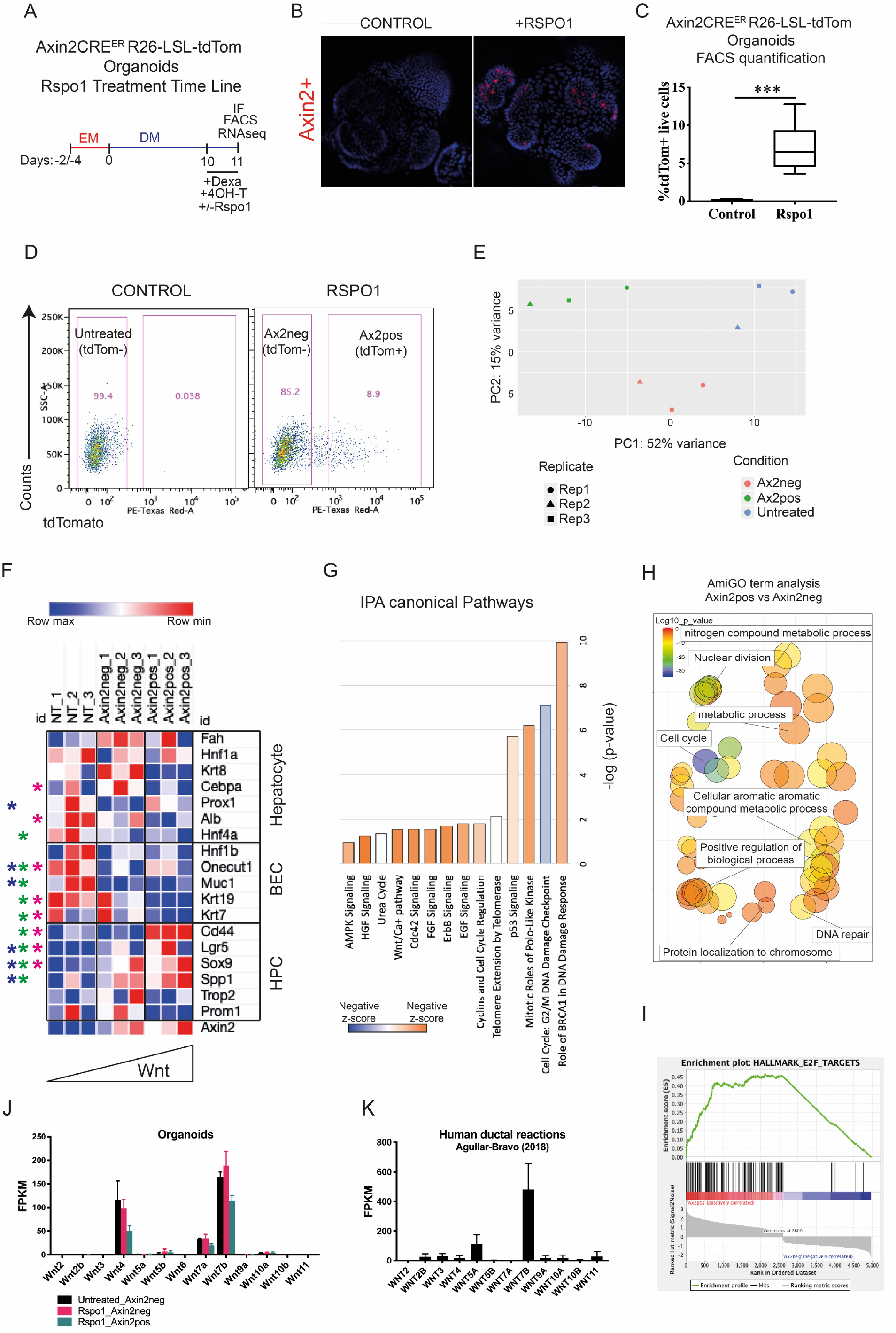
(A) Treatment timeline of Axin2CreERT2 organoids for panels D to J. (B) Representative IF images of control and Rspo1 treated organoids (see panel A). In red, endogenous tdTom labels Axin2pos subpopulation. Cell nuclei counterstained with DAPI (blue). Scale bar 100 μm. (C) Bar graph shows flow cytometry quantifications for the percentage of Axin2pos cells in differentiated organoids (see panel A) following 24h 4-OHT Rspo1 treatment. n=7 (organoids derived from 3 animals were analysed at 7 different passages using 6 different differentiation batches (Table S2)) (D) FACs gating strategy used for RNAseq. Flow cytometry dot plots show the presence of Axin2pos in differentiated organoids treated with Rspo1 but not in control conditions (see panel A). (E) PCA of Axin2pos, Axin2neg and untreated (NT) cells showed clustering of the samples based on treatments. (F) RNAseq FPKM gene expression values of various hepatocyte, HPC and BEC lineage markers. Blue asterisk indicates differentially expressed (padj≤0.01) between untreated and Axin2neg cells. Green asterisk indicates genes differentially expressed (padj < 0.01) between Axin2neg and Axin2pos cells. Magenta asterisk indicates genes differentially expressed (padj<0.01) between untreated and Axin2pos cells. (G) IPA canonical pathway analysis of 2-fold change differentially expressed genes (padj value <0.01) between Axin2^pos^ and Axin2^neg^ cells (1164 genes). (H) REVIGO scatterplot of the enriched GO terms from Rspo1-treated BD organoids RNAseq experiment. GO enrichment analysis for top most variable 500 transcripts in PC1 when comparing Axin2pos vs Axin2neg PC1. Colors reflect the padj associated with the GO term categories while circle size reflects the number of genes relating to the term. (I) GSEA analysis using MSigDB C2 comprising Curated Gene Sets from online pathway databases publications in Pubmed and knowledge of domain experts reveal enrichment of E2F target genes in Axin2pos cells when compared to Axin2neg cells. Analysis was performed in genes differentially expressed genes between Axin2pos and Axin2neg with a padj < 0.01. A total of 5333 genes where found differentially expressed. Of those 5047 had a human orthologue and therefore could be analysed using GSEA. (J) FPKM gene expression values according to RNAseq of untreated, Axin2pos, Axin2neg. (K) FPKM gene expression values of CK7+ regions from LSC RNAseq experiments of Aguilar-Bravo et al (2018)

In Rspo1-treated organoids two cell populations (Axin2^pos^ and Axin2^neg^) could be distinguished based on tdTom expression (Figure 4, D). We next uncovered the transcriptome profiles of these organoid cell subpopulations by subjecting sorted Axin2^pos^ and Axin2^neg^ cells to RNAseq analysis. Additionally, we sequenced cells from untreated organoids which were all negative for tdTom expression (Figure 4, D and Figure S1, B). Principal component analysis (PCA) showed separation of the three sequenced populations based on treatments (Figure 4, E). When comparing hepatocyte, BEC and HPC markers, the biggest differences in gene expression profiles were observed between Rspo-treated and untreated cell populations, suggesting that cells sorted as Axin2^neg^ also responded to the Rspo1 to a certain extent (Figure 4, F). The addition of Rspo1 to the cultures caused an overall decrease of BEC lineage markers (Hnf1b, Muc1, Krt19, Krt7) in both Axin2^pos^ and Axin2^neg^ cells when compared to untreated cells, indicating that exposure to Rspo1 promoted escape from biliary fate (Figure 4, F). By contrast, the expression of hepatocyte lineage markers in Rspo1 treated cells did not follow a consistent trend, with only Hnf4a displaying a decrease when comparing Rspo1-treated (Axin2pos and Axin2neg) and untreated cells (Figure 4, F). The expression of Lgr5, Sox9 and Spp1 was significantly increased in Rspo1-treated Axin2^pos^ and Axin2^neg^ cells when compared to untreated cells, indicating that some but not all HPC markers were increased in response to 24h Rspo1 treatment (Figure 4, F). Overall, these results support the idea that activation of Wnt signalling in BD organoids promotes both an escape from biliary fate and an increase in the expression levels of HPC markers, while preventing the acquisition of the hepatocyte lineage marker Hnf4a.

PCA revealed differences in the transcriptome of Axin2^pos^ and Axin2^neg^ cells. To further investigate such differences, we subjected Axin2^pos^ and Axin2^neg^ cell subpopulations to differential expression analysis. Ingenuity Pathway Analysis (IPA) revealed upregulation of canonical pathways in Axin2^pos^ cells including “Brca1 DNA damage responses”, “Cell cycle and G2/M DNA damage checkpoint”, “mitotic roles of Polo-like kinase and DNA double-strand break repair by homologous recombination” (Figure 4, G). Similarly, AmiGO Gene Ontology (GO) analysis of the top 500 of PC1 were enriched in GO terms related to cell division, including “Cell cycle”, “Nuclear division”, “DNA repair” and “Protein localization to chromosome” (Figure 4, H). Gene Set Enrichment Analysis (GSEA) further showed enrichment of E2F target genes which regulate G1/S transition in Axin2^pos^ cells (Figure 4, I). These results are therefore consistent with the recent observations by Spit et al. (2020), who showed that activation of Wnt/Rspo signaling though the expression of an oncogenic form of RNF43 triggered the activation of DNA damage response and cell division transcriptional signatures in colon epithelial cells (Spit et al., 2020).

Interestingly, IPA analysis showed that Axin2^pos^ cells were enriched in genes involved in the “Wnt/Ca2+ pathway”, raising the possibility that in addition to the Wnt/β-catenin pathway other branches of the Wnt signaling cascade might have been activated in response to Rspo1 treatment (Figure 4, G). Consistently, BD organoid cells were found to express various Wnt co-receptors involved in the transduction of β-catenin-independent intracellular cascades (Ryk, Ptk7, Vangl1 and to a lesser degree Rora, Ror1, Vangl2, Prickle1, Prickle3 and Prickle4) (Figure S4, C-F). BD organoid cells were also expressed various Wnt ligands with described canonical and non-canonical activity, with Wnt7b, Wnt7a and Wnt4 the most abundantly expressed (Figure 4, J) (Eubelen et al., 2018; Vallon et al., 2018; Yu et al., 2008) (Alok et al., 2017; Cho et al., 2017; Ferrari et al., 2018; Le Grand et al., 2009; Posokhova et al., 2015). To assess whether the panel of Wnts expressed by BD organoids might be relevant in a clinical setting, we examined the expression of Wnt ligands in the ductal reactions of human cirrhotic patients using the public data from laser capture microdissection (LCM) experiments (Aguilar Bravo et al., 2019). Wnt7b was the most abundantly expressed Wnt ligand in ductal reactions of cirrhotic human samples dissected based on CK7+ expression, suggesting that the biliary epithelium might be an important contributing source to the ductal reactions Wnt microenvironment of these patients (Figure 4, K).

In the organoid system, Axin2^pos^ cells had enhanced expression of HPC markers when compared to Axin2^neg^ cells. To explore whether these transcriptional differences had a functional impact in the progenitor properties of the cells, the ability of Axin2^pos^ and Axin2^neg^ cells to form secondary organoids was evaluated. Axin2^pos^ cells showed significantly higher organoid forming capacity when compared with the Axin2^neg^ cell population of Rspo1-treated organoids (Figure 5). Altogether, these data indicate that differentiated BD organoids are a source of Wnt ligands that might be relevant in a clinical setting and that these in combination with Rspo ligands likely induce the appearance of an Axin2 expressing population with progenitor features.

**Figure 5.**
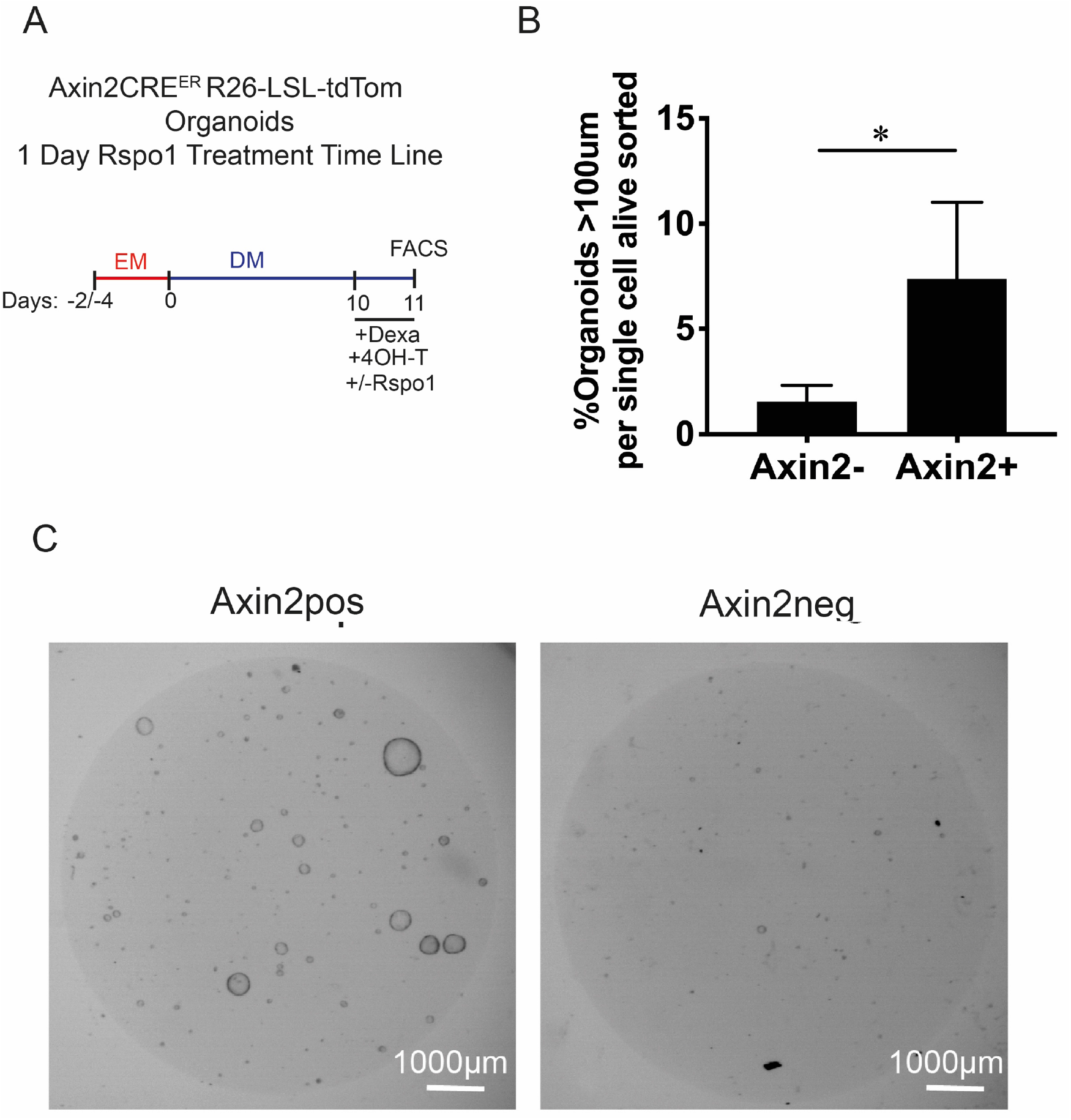
(A) Treatment timeline of Axin2CreERT2 organoids for panels B and C.(B) Bar graph shows organoid forming capacity of Axin2pos and Axin2neg sorted cells from Axin2CreERT differentiated BD organoids exposed to Rspo1 for 1 day. Panel E indicates experimental design. n= 5 (organoids derived from 2 animals were analysed at 5 different passages using 5 different differentiation batches (Table S2)). SD error bars. Statistics were determined using paired t-test. (C) Representative images for panel B quantifications showing organoids formed from Axin2pos and Axin2neg sorted cells (333 cells sorted cells per 50μl Matrigel) after 7 days. Axin2pos cells formed more organoids than Axin2neg cells.

## Discussion

Here we report that the sole addition of Rspo1 in BD organoid cultures was sufficient to induce the appearance of an organoid cell population with progenitor features characterized by high canonical Wnt activation levels and enhanced organoid forming capacity. Organoids primarily expressed Wnt4, Wnt7b and Wnt7a and Rspo1 HPC-induced signature was dependent on these endogenously produced Wnt ligands. Of the organoid expressed ligands, Wnt7b was strongly expressed in DR of human cirrhotic patients (Aguilar Bravo et al., 2019). A range of other studies also demonstrated expression of Wnt7a and Wnt7b ligands in BECs following liver injury (Okabe et al., 2016; Pepe-Mooney et al., 2019);(Boulter et al., 2015). Together, these observations support that the biliary epithelium is an important contributing source of Wnt ligands in liver disease and that Rspo ligands act as the limiting factor controlling the activation of canonical Wnt pathway in the biliary epithelium and exit from biliary fate following severe liver damage. The expression of Rspo family members was not detected in BD organoids, suggesting that other cell types may be responsible for its presence in pathophysiological contexts *in vivo*.

Wilson et al. (2020) recently reported that Axin2 transcripts were detectable in homeostatic BECs, but that levels did not increase during DDC peak injury using a liver damage strategy that did not force BEC-to-hepatocyte conversion (Wilson et al., 2020). Here we show that the Wnt/β-catenin pathway is dynamically activated in BECs isolated from AAV8.p21 but not from AAV8.null MCD injured livers following withdrawal of the hepatotoxic insult, suggesting that the biliary epithelium is able to respond to canonical cues when hepatocyte proliferation is compromised and that this is an event associated with BEC reprograming. Our data is also consistent with RNAseq pathway enrichment analysis performed in chronically injured human cirrhotic livers which also identified activation of the Wnt/β-catenin pathway in BECs (Spee et al., 2010) (Ceulemans et al., 2017). Wilson et al. (2020) and our observations however contrast with the studies of Planas-Paz et al. (2019) and Pepe-Mooney et el. (2019), that failed to identify Axin2 expression in BECs isolated from DDC-injured livers using 10x genomics single-cell sequencing (scRNAseq) (Pepe-Mooney et al., 2019; Planas-Paz et al., 2019). We attribute these discrepancies to differences in the sensitivities of the 10x genomics scRNAseq and RT-qPCR techniques.

Axin2 expression was most strongly upregulated in the biliary epithelium at day 6 of recovery of AAV8.p21 MCD injured livers. Interestingly, experiments performed by Russell et al. (2019) showed that BEC-to-hepatocyte conversion in β-catenin KO MCD-injured livers primarily occurred between day 3 and day 7 recovery, which is consistent with the idea that strongest activation of the Wnt/β-catenin in the biliary epithelium coincides with point at which the number of cells transitioning towards hepatocyte fate is maximal (Figure 6) (Russell et al., 2019). Previous studies have shown that activation of Wnt/β-catenin signalling was critical for the generation of proliferative bipotent hepatoblasts during the differentiation of ESCs into hepatocytes (Touboul et al., 2016). These data, together with our observations in the organoid system supports the hypothesis that activation of the Wnt/β-catenin pathway promotes the acquisition of a HPC-like transient state. Of note, activation stabilization of β-catenin prevented the expression of Hnf4a. It is therefore possible that downregulation of Wnt/β-catenin pathway is necessary for bi-potent HPCs to progress towards hepatocyte fate.

**Figure 6.**
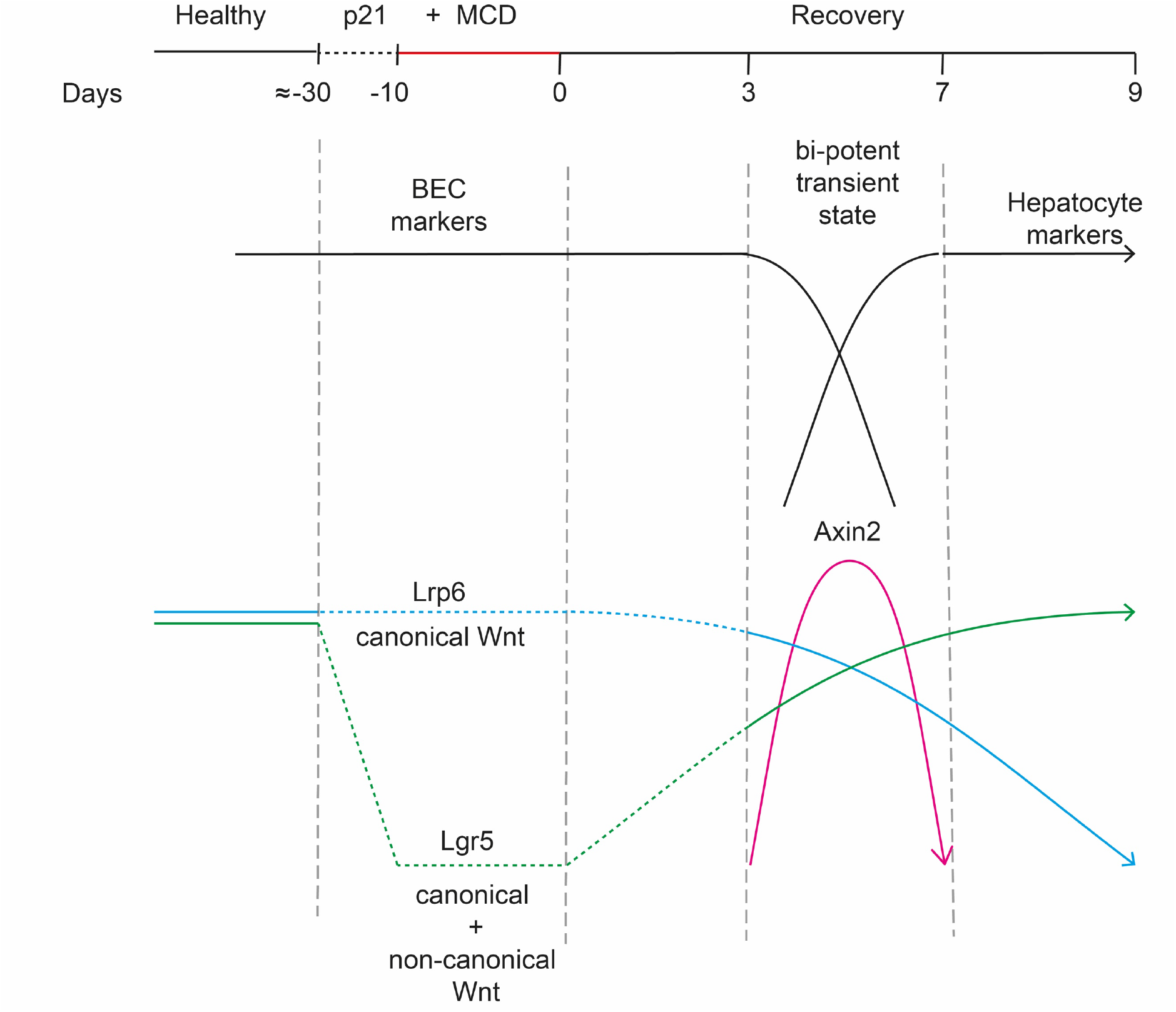
Scheme integrating the dynamics of BEC-to-hepatocyte transdifferentation and the expression patterns of Axin2, Lrp6 and Lgr5 obtained by RT-qPCR analysis of BECs (tdTom+ Epcam+) isolated at different time points during the recovery period of the p21 MCD mouse model. BEC-to-hepatocyte conversion occurs predominantly between day 3 and day 7 of the recovery period (Russell et al., 2019). Lrp6 gene expression (blue line) gradually declines between day 3 and day 9. By contrast, Lgr5 gene expression (green line) gradually increases between day 3 and day 9 of the recovery period. Expression of the Wnt/β-catenin target gene Axin2 (magenta line) is significantly increased at day 6 of recovery. Sections represented with dash lines represent predicted gene expression levels.

Under prolonged or severe damage, hepatocyte proliferation is compromised which translates in deficient liver regeneration and causes progressive loss of organ function. Given that hepatocyte proliferation in injured livers is dependent on canonical Wnt cues, the pharmacological activation of the Wnt/β-catenin pathway may seem a priori an attractive therapy to boost the hepatocyte-driven repopulation. However, the data presented here suggest that activation of Wnt signalling in patients suffering from chronic liver damage may prevent hepatocyte and biliary differentiation from cells in an intermediate differentiation state. In fact, experiments carried out by the Boulter and Kohara laboratories have shown that pharmacological depletion of Wnt secretion in murine livers promotes resolution of DR and ameliorated hepatitis C virus- and DDC-induced fibrotic scarring, suggesting that systemic inhibition of the Wnt pathway may be an attractive therapy for patients suffering from chronic liver disease (Tokunaga et al., 2017) (Wilson et al., 2020). A better understanding roles and timing of Wnt signalling processes during chronic liver damage that unravels the complex cellular communication *in vivo* should help in the design of treatments that can be tailored to the pathological setting of each individual patient.

## Experimental Procedures

### Animal Experimental Models

All animal work was carried out under UK Home Office project and personal licenses following local ethical approval and in accordance with local and national guidelines. Axin2-CreERT2 mice were a gift from Chambon laboratory and were crossed with R26-LSL-tdTom mice (The Jackson Laboratory). Tet-O-ΔN89β-catenin animals were generated in house (Jardé et al., 2012). Experiments with Krt19-CreERT2; R26-LSL-tdTom animals were conducted at the Centre for Regenerative Medicine, The University of Edinburgh. 8-week old animals were injected intraperitoneally with 3 doses of 4 mg tamoxifen (Sigma) and allowed a 2-week wash-out period. Animals received a single dose of AAV8.TBG.null or AAV8.TBG.p21 (5*10^12 units/ml) (Addgene) intravenously allowed 1 week wash-out period on normal diet. The mice were fed MCD diet (Dyets Inc.) for 15 days and then switched to normal mouse chow for the recovery period. Animals were sacrificed via a Schedule 1 method on recovery days 3, 6 and 9 and the liver was extracted for downstream analysis.

### Generation and differentiation of liver organoids of biliary duct origin

Liver organoids were generated from BD isolated as previously described (Huch et al., 2013). BD organoid expansion medium was defined as AdDMEM/F12 containing 1% B27 supplement, 0.5% N2 supplement, 1.25 μM N-Acetylcysteine, 10 nM gastrin, 100 ng/ml FGF10, 50 ng/ml EGF, 50 ng/ml HGF, 10 nM Nicotinamide and 10% Rspo1 Conditioned Medium). For the first 7 days of culture after isolation, expansion medium was supplemented with 25 ng/ml Noggin. For differentiation, organoids were expanded for 2 to 4 days, and switched to differentiation media (AdDMEM/F12 containing 1% B27 supplement, 0.5% N2 supplement, 1.25 μM N-Acetylcysteine, 50 ng/ml EGF, 100 ng/ml FGF10, 50nM A-8301, 10 μM DAPT for 13 days. Organoids were routinely 1:5 passaged by mechanical disruption every 5 to 8 days. When seeded for experiment, organoids were incubated with TriplE (Gibco) for 7min at 37C, digested into single cells and seeded at a concentration of 200-400 cells/μl matrigel. Differentiation media was changed every day and supplemented with 3 μM dexamethasone for the last 3 days of culture. When indicated, differentiated organoids were treated with Rspo1 (100 ng/ml in 0.1% BSA PBS), doxycycline (0.1 mg/ml in PBS), 4-OHT (500 nM in ethanol), Wnt3a (100 ng/ml in 0.1% BSA PBS) or LGK974 (500 nM in DMSO). Organoids used in this study were between passage 4 and 15.

### Isolation of Axin2^pos^ and Ax2^neg^ cells by flow cytometry

Organoids were collected, dissociated into single cells (7min at 37°C with TriplE) and sorted into 0.5 ml of expansion media or 0.5 ml of Trizol for organoid forming capacity assay or RNA sequencing, respectively. FACS gating strategy for Axin2^pos/neg^ isolation is described in Figure S1, A.

### Isolation of tdTom+ Epcam+ biliary cells from MCD-injured livers

Liver tissue (cut into ~10mm^2^ cubes) was digested at 37° C in collagenase XI and dispase II dissolved in DMEM/Glutamax at concentration 0.125 mg/ml. Following digestion, cells were washed and dissociated into single cells at 37° C in TrypLE. Cells were stained with antibodies for Epcam (G8.8; eBiosciences), CD45 (30-F11; Biolegend), CD31 (390; Biolegend) and Ter119 (TER119; Biolegend). Dead cells were excluded using DAPI. tdTom+/Epcam+/CD45-/CD31-/Ter119-cells were isolated using BD FACS Fusion at the Flow and Genomics Facility, Centre for Regenerative Medicine, The University of Edinburgh.

### Organoid forming capacity assay

Sorted cells were plated in expansion media containing 25 ng/ml Noggin with 10% Rspo1 CM or 300 ng/ml of purified Rspo1 as indicated. 10 μM Y27632 was added for the 3 first days of culture. At day 7, images were acquired using a GelCount instrument and organoid diameter was quantified with ImageJ. Structures with a diameter >100 μm were considered as organoids for the analysis.

### RNA isolation and gene expression analysis

RNA from tdTom+ Epcam+ cells from MCD-injured liver isolated as before was extracted using PicoPure RNA isolation kit (Thermo Fisher). RNA from primary hepatocytes and organoids was isolated following Trizol Reagent RNA extraction method coupled with RNaesy MinElute columns (Qiagen). cDNA was synthetized using an iScript cDNA Synthesis Kit from Bio-Rad, subsequently diluted in water (1:4) and mixed with SensiFAST SYBR Hi-Rox (Bioline) and the appropriate primers at a final concentration of 10 μM. Three technical replicates were performed per condition. Gene expression was normalized to B2M. The primer pairs used in this study are listed in Table S1.

### Immunofluorescence

Organoids were fixed either with 4% PFA for 1h at RT or with ice-cold 100% methanol for 30 min at −20°C, washed 3 times with PBS and permeabilized/blocked for 3-5h at RT with 10% donkey serum in IF buffer (0.1% Triton X-100, 0.05% Tween-20, 0.1% BSA in PBS). Primary antibody incubation was performed in IF buffer overnight at 4° C. Cells were then washed (IF buffer), incubated with secondary antibody for 3-5h at RT, washed (IF buffer) and counterstained with DAPI in PBS for 15min. Samples were imaged in PBS. Primary antibodies used were as follow: anti-HNF4a (PP-H1415-00, R&D systems), anti-RFP (600-401-379, Rockland), anti-CK19 (TROMA-III, DSHB).

### Organoid viability assay

Organoid viability was the determined by CellTiter-Glo (Promega) and CellTox (Promega) following manufacturer’s instructions.

### RNAseq of Axin2^pos/neg^ and untreated organoid cells

Total RNA of Axin2^pos/neg^ sorted cells was isolated as above (Figure S1,B) from organoids derived from the same animal were analysed at 3 different passages using 3 different differentiation batches (n=3) (Table S2). Single-end 75bp sequencing was performed with 100ng of total RNA input on a NextSeq-500 sequencer where equimolar libraries were pooled. Sequences were trimmed (Trimmomatic 0.35), mapped to the mouse GRCm38 reference genome (STAR 2.5.1b) and counts were assigned using featureCounts (Dobin et al., 2012) (bolger et al., 2014; Liao et al., 2014). Genes with an average of fragments per kilobase per million (FPKM) <0.5 in all biological groups were considered as not expressed and discarded from the analysis. Differential gene expression was performed using DESeq2 (Love et al., 2014). Data have been deposited as GSE161039 in the Gene Expression Omnibus (GEO) database.

### Statistical analysis

GraphPad Prism 7.0 Software was used for all statistical analyses. Statistics of RT-qPCR analysis correspond to a p-value n.s.>0.05; p-value *< 0.05; p-value **< 0.01; p-value ***< 0.001; p-value ****< 0.0001.

## Acknowledgements

We thank Dr Alexander Raven for helpful discussions. We thank Mark Birshop and Jolene Twomey for assistance with FACS, Angela M. Marchbank and Georgina E. Smethurst for the RNAseq library preparation and Robert Andrews and Sumukh Deshpande for assistance with the bioinformatic analysis. This work was supported by the Marie Sklodowska-Curie Actions Innovative Training Network 675407 (acronym PolarNet) and the William Morgan Thomas Fund. MM, and TGB were funded by the Wellcome Trust (Grant number: WT107492Z).

## Author Contributions

E.V.E. conceived the study and drafted the work; E.V.E., N.A., C.V.M., M.D., M.Z. performed the experiments; M.M. and T.G.B. provided essential tools; E.V.E. and N.A. analysed and interpreted the data; E.V.E., N.A., M.M., A.O., M.J.S., C.H., S.J.F., T.G.B and T.C.D. made important intellectual contributions; all authors contributed to finalizing the manuscript.

## Supplemental Information

**Figure S1.**
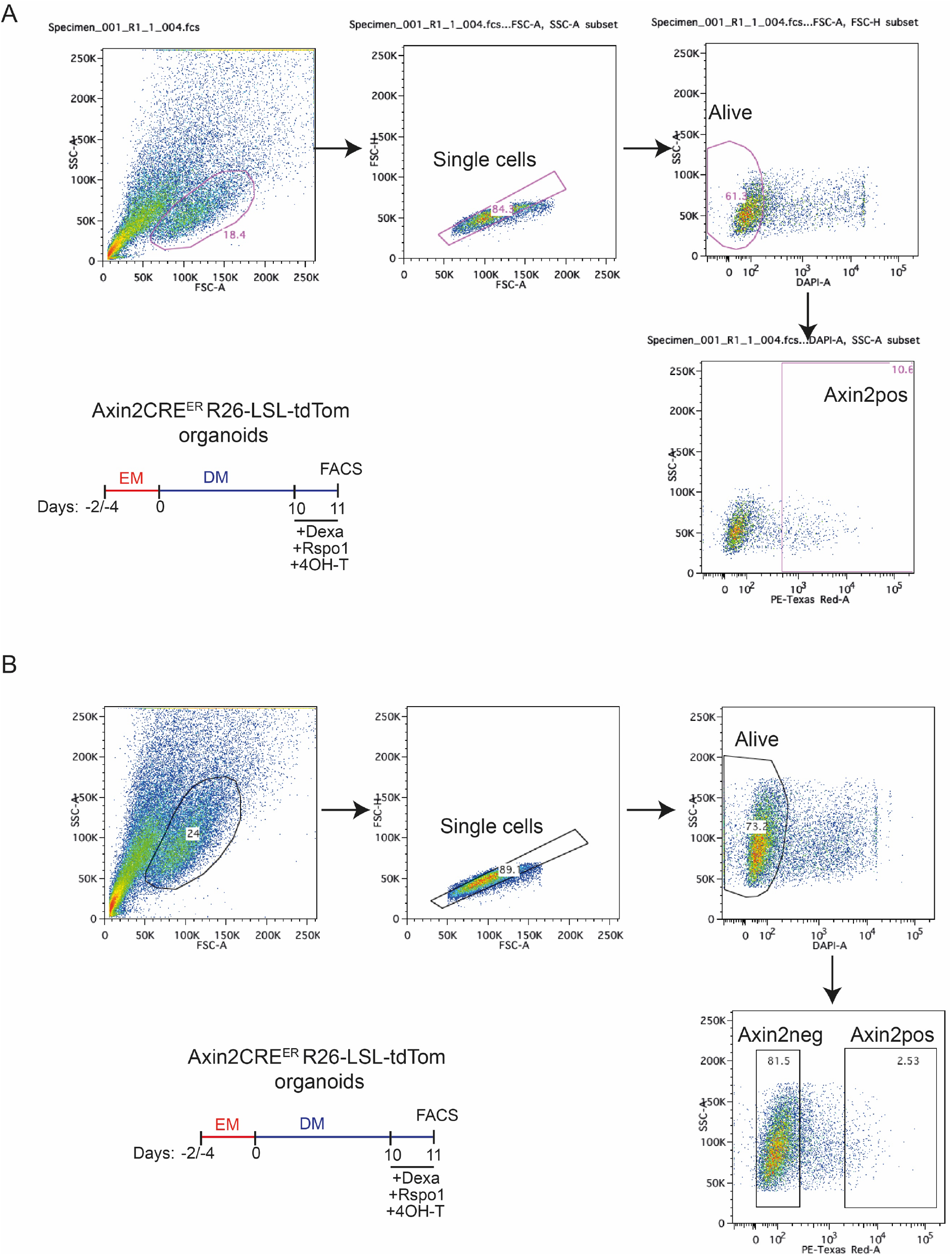
(A) FACS gating strategy for the quantifications of Axin2 expressing cells positives for tdTom (B) FACS gating strategy for the isolation of Axin2pos and Axin2neg cells isolated from the AxineCreERT2; R26-SLS-tdTom mouse model.

**Figure S2.**
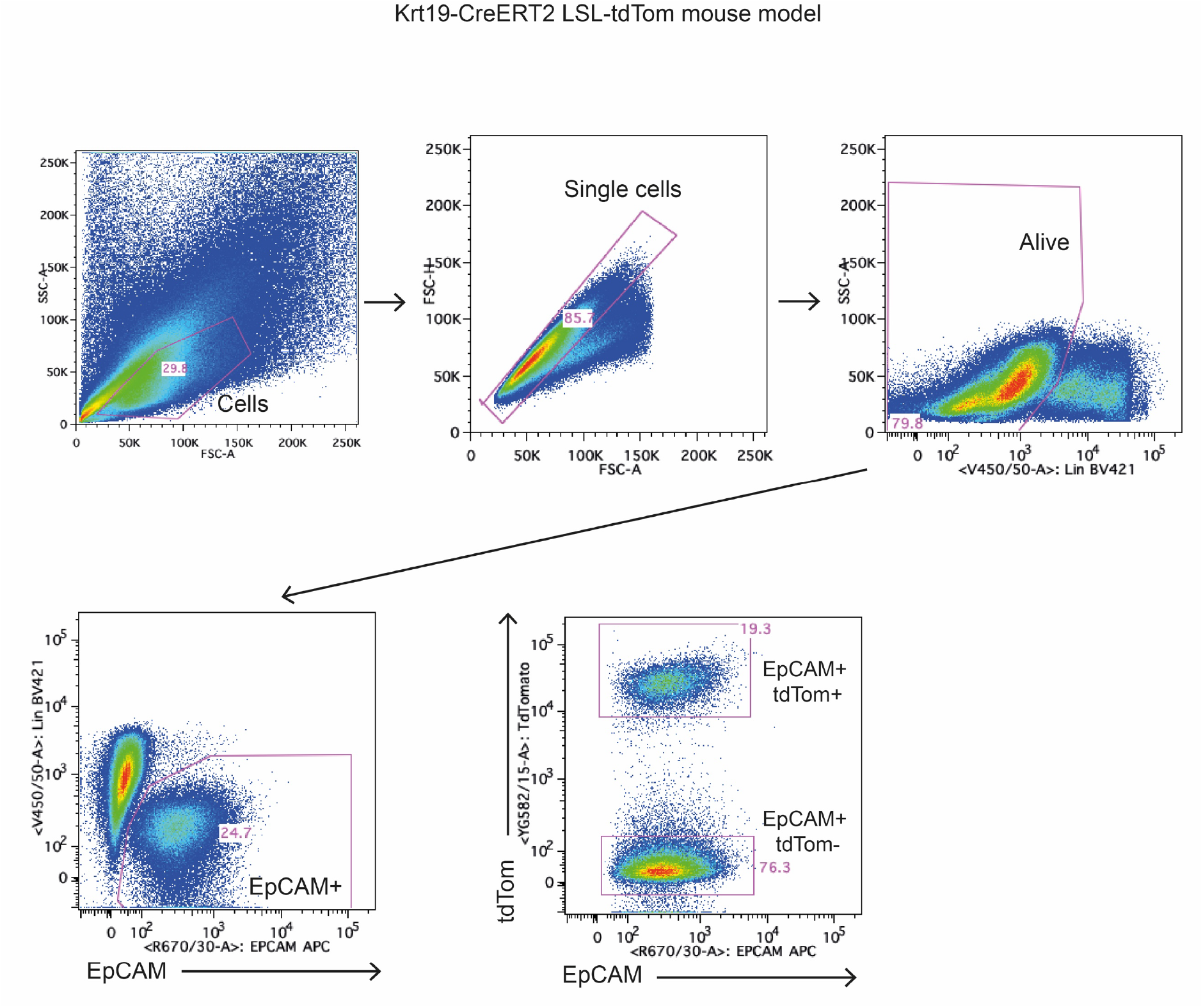
FACS gating strategy for the isolation of BECs in the Krt19-CreERT2; R26-LSL-tdTom following tamoxifen administration and at different times points MCD liver injury recovery.

**Figure S3.**
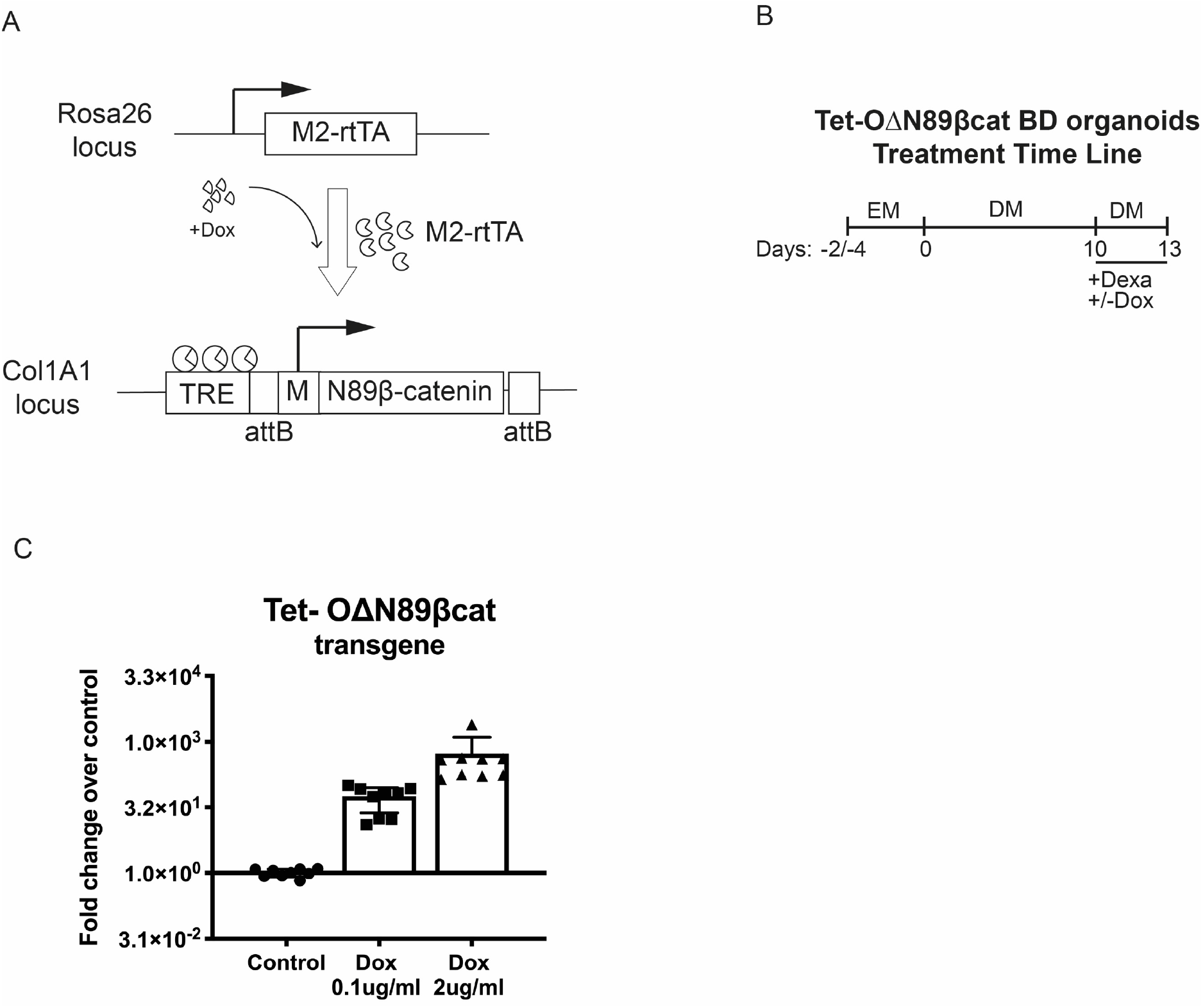
(A) Schematic representation of Tet-OΔN89β-catenin and M2-rtTA transgenes present in Tet-OΔN89β-catenin organoids. Presence of doxycyclin (Dox) triggers the transcription of activated (ΔN89) mutant β-catenin. (B) Treatment timeline for panel C. (C) RT-qPCR shows expression of the OΔN89β-catenin transgene upon doxycycline treatment. n=9 (organoids derived from a single animal were analysed at 3 different passages using 3 different differentiation batches. In each passage, organoids were cultured in 3 wells that were processed independently (Table S2)). SD error bars. Statistics were determined using multiple t-test.

**Figure S4.**
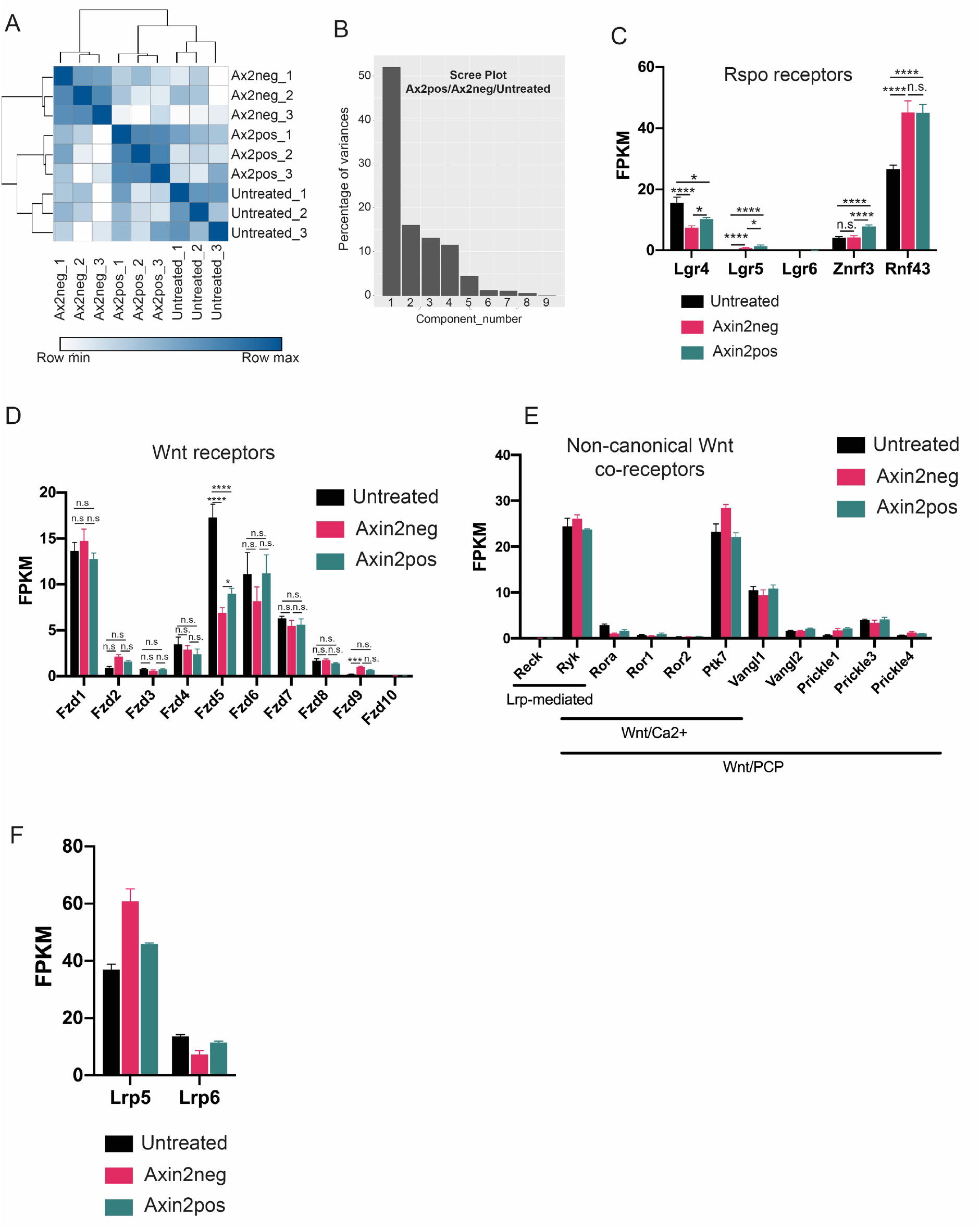
(A) Heatmap showing hierarchical clustering of Axin2pos, Axin2neg and untreated cells based on sample distances calculated with R. (B) Scree plot showing the eigenvalues for each of the dimensions of the PCA. (D-F) Expression levels of Rspo and Wnt related receptors in Axin2pos, Axin2neg and untreated cells by RNAseq expressed in normalized FPKM values.

**Table S1.**
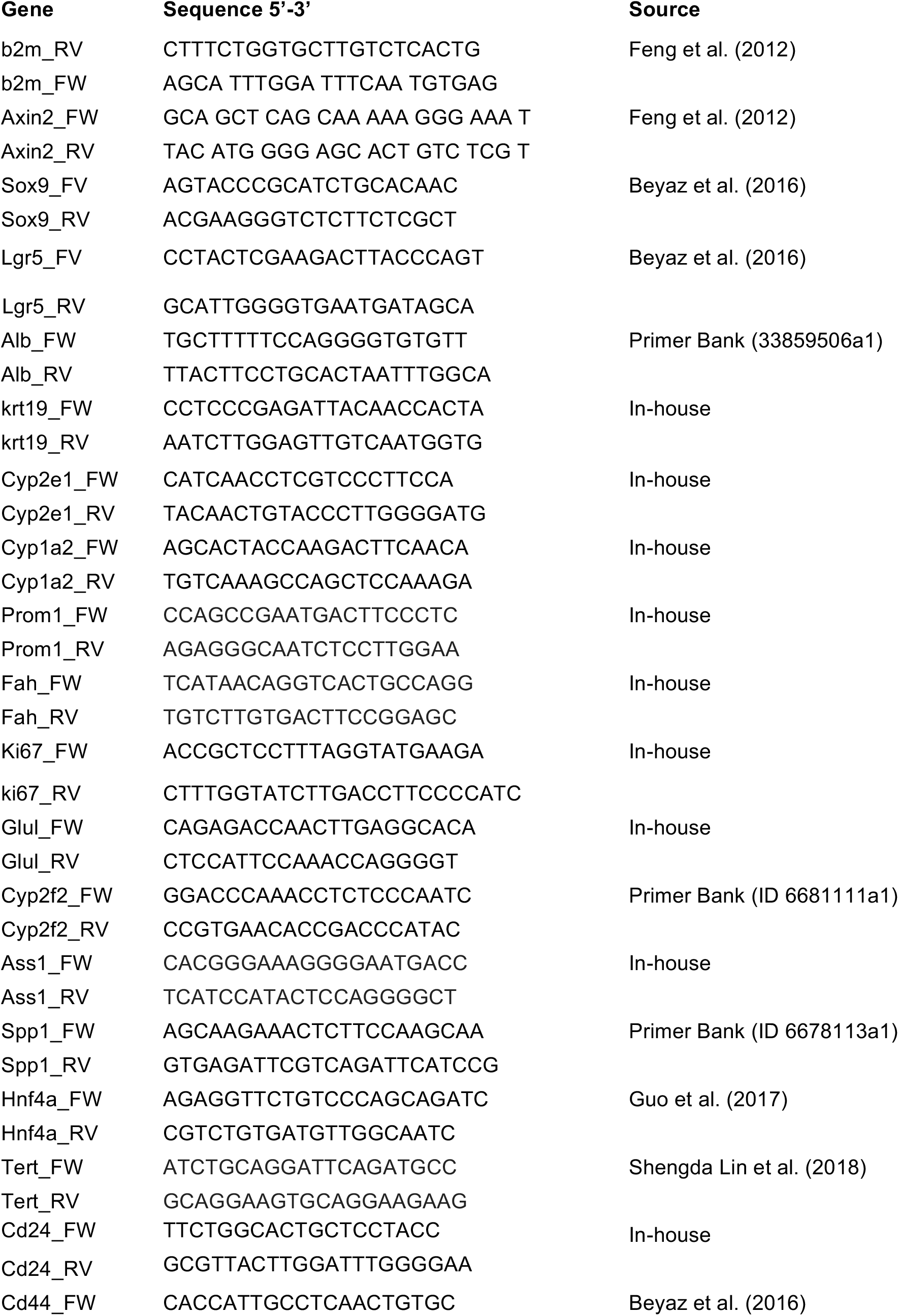
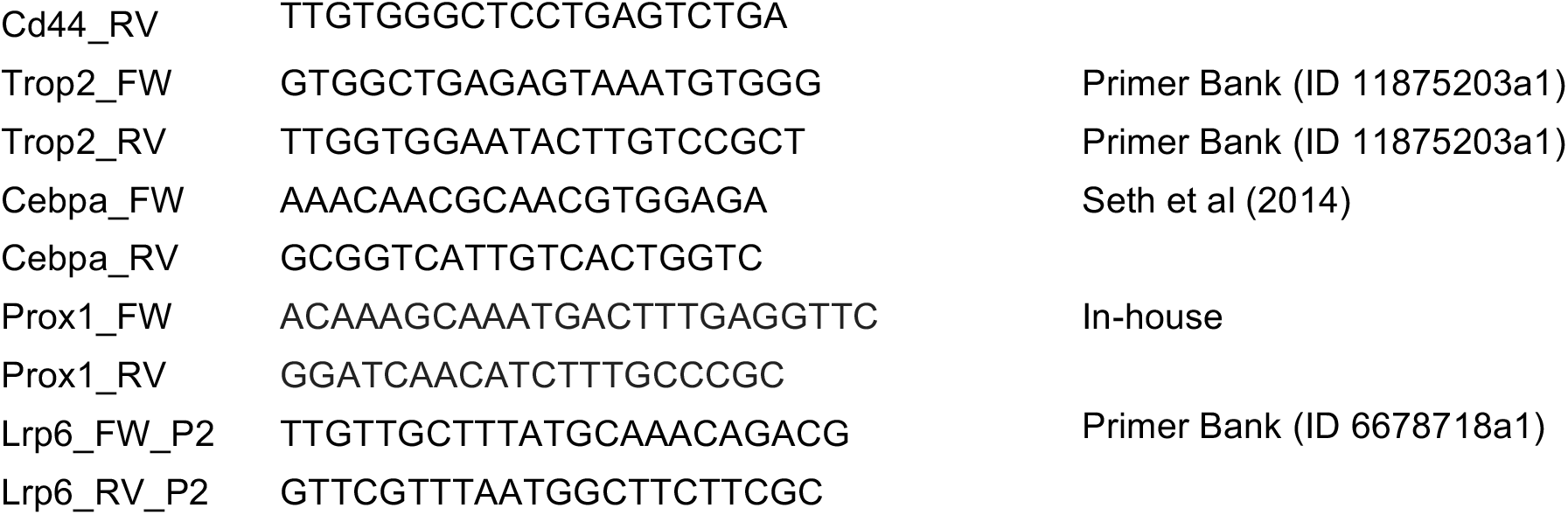
RT-qPCR primer pairs used in this study.

**Table S2.** Organoid lines used in this study

| Organoid ID | Information animal of origin |  |  | Passage | Differentiation Batch | Independent wells* | Data |
| --- | --- | --- | --- | --- | --- | --- | --- |
|  | Genotype | Sex | Genetic background |  |  |  |  |
| 1115 | WT | M | C57BL/6J | P5 | A | 3 | Fig2 (B-D) |
| 1106 | WT | M | C57BL/6J | P4 | B | 3 | Fig2 (B-D) |
| 1119 | WT | M | C57BL/6J | P7 | C | 3 | Fig2 (B-D) |
| Tet-O | Tet-O<br>ΔN89βcat | M | Mixed | P8 | D | 3 | Fig3(B,D,E)<br>S3 |
| Tet-O | Tet-O<br>ΔN89βcat | M | Mixed | P10 | E | 3 | Fig3(B,D,E)<br>S3 |
| Tet-O | Tet-O<br>ΔN89βcat | M | Mixed | P13 | F | 3 | Fig3(B,D,E)<br>S3 |
| BAL811 | WT | M | BALB/cJ | P8 | G | 1 | Fig3, C |
| BAL211 | WT | M | BALB/cJ | P12 | H | 1 | Fig3, C |
| Balbc | WT | M | BALB/cJ | P4 | I | 1 | Fig3, C |
| T124 | WT | M | Mixed | P16 | I | 1 | Fig3, C |
| T124 | WT | M | Mixed | P17 | J | 1 | Fig3, C |
| RMD21.4b | Ax2Cre<sup>ER</sup><br>R26-LSL-tdTom | M | Mixed | P5 | K | 1 | Fig4, C<br>RNAseq |
| RMD21.4b | Ax2Cre<sup>ER</sup><br>R26-LSL-tdTom | M | Mixed | P6 | L | 1 | Fig4, C<br>RNAseq |
| RMD21.4b | Ax2Cre<sup>ER</sup><br>R26-LSL-tdTom | M | Mixed | P7 | M | 1 | Fig4, C |
| RMD21.4b | Ax2Cre<sup>ER</sup><br>R26-LSL-tdTom | M | Mixed | P8 | N | 1 | Fig4, C |
| RMD21.4b | Ax2Cre<sup>ER</sup><br>R26-LSL-tdTom | M | Mixed | P10 | O | 1 | Fig4, C<br>RNAseq |
| RMD24.2b | Ax2Cre<sup>ER</sup><br>R26-LSL-tdTom | M | Mixed | P12 | P | 1 | Fig4, C |
| RMD22.2g | Ax2Cre<sup>ER</sup><br>R26-LSL-tdTom | F | Mixed | P9 | Q | 1 | Fig4, C |
| RMD21.4b | Ax2Cre<sup>ER</sup><br>R26-LSL-tdTom | M | Mixed | P5 | K | 3 | Fig5 |
| RMD21.4b | Ax2Cre<sup>ER</sup><br>R26-LSL-tdTom | M | Mixed | P6 | L | 3 | Fig5 |
| RMD21.4b | Ax2Cre<sup>ER</sup><br>R26-LSL-tdTom | M | Mixed | P7 | M | 3 | Fig5 |
| RMD21.4b | Ax2Cre<sup>ER</sup><br>R26-LSL-tdTom | M | Mixed | P10 | O | 3 | Fig5 |
| RMD24.2b | Ax2Cre<sup>ER</sup><br>R26-LSL-tdTom | M | Mixed | P12 | P | 3 | Fig5 |

\*Organoids from the same organoid line at the same passage were cultured in parallel in different wells. Biological samples from these wells were not mixed and processed independently for downstream proposes.

